# Strategies of mechanical adaptations of circulating tumor cells and properties of their cellular microenvironment have predictive and prognostic biomarker value

**DOI:** 10.64898/2026.10.31.763352

**Authors:** Maria Gaczynska, Fatemeh Ataei, Matthew Alan Click, Chih-Wei Chou, Furkan Dursun, Chia-Nung Hung, Kerri A. Kendrick, Robin Leach, Michael Liss, Penny Vroman, Zhao Zhang, Tim H.-. Huang, Pawel A. Osmulski

## Abstract

Mechanically demanding conditions in the blood circulation damage and kill majority of tumor-shed circulating tumor cells (CTCs). Mechanical fitness: a set of mechanical properties detected with atomic force spectroscopy based force spectrometry supports survival of CTCs. We propose that live CTCs isolated from peripheral blood of prostate cancer patients represent distinct strategies of fitness promoting mechanical adaptations to the fluid shear stress. Some of the cells mimicked properties of erythrocytes (high stiffness) or leukocytes (high deformability). The clustering strategy characterized by exceptionally high adhesion is ingenious to CTCs and not found in blood cells. The abundance of CTCs with elevated adhesion has a predictive value for prostate cancer patients starting 1^st^ round of androgen deprivation therapy. Moreover, a “high adhesion score” based on individual cells’ or average patient-specific adhesion is instrumental to stratify recurrent patients that underwent diagnostic imaging by PSMA-PET/CT and clarify inconclusive PSMA scan results. Including contribution of erythrocyte-like strategy broadened the predictive power of CTC mechanophenotyping. Improved biomarking opportunities to predict early castration resistances were noted when the composition of CTC microenvironment was included alongside the mechanical properties. We demonstrated previously that the CTC microenvironment is formed by large cells co-purifying by microfiltration of the blood of cancer patients. Analysis of the association of particular components of CTC microenvironment with the clustering strategy led to pointing at highly adhesive CTC-EMT. However, in the advanced patients adhesiveness of CTC-EMT was surpassed by tumor hybrid cells (THCs), a distinctive kind of invasive species formed by incomplete phagocytosis of CTCs by macrophages. THCs changed their mechanical phenotype and strategy preference with disease progression pointing at the dynamic nature of adaptations of tumor-shed cells to circulation conditions.

## Introduction

Circulating tumor cells (CTCs) are shed into the bloodstream from epithelial-origin tumors and active cancer lesions, where they can travel to distant tissues, extravasate, and initiate metastatic growth¹–³. Initially characterized by epithelial features, including specific surface markers and relatively large cell size, CTCs can be isolated as viable cells from liquid biopsies obtained from just a few milliliters of peripheral blood. In prostate cancer (PC), CTCs are detected at higher numbers in patients with high-risk or metastatic disease, and their abundance generally increases with disease progression⁴. Prostate cancer is typically a slowly progressing disease, and progression from localized disease to metastasis may take years or even decades⁵. Following initial diagnosis and treatment, often involving surgical removal of the primary tumor, the disease may remain clinically dormant for extended periods. However, an increasing level of prostate-specific antigen (PSA) can indicate disease recurrence, referred to as biochemical recurrence (BCR). In current clinical practice, patients with BCR frequently undergo prostate-specific membrane antigen positron emission tomography/computed tomography (PSMA-PET/CT) to detect local or distant disease recurrence. Although PSMA-PET/CT can identify the extent and location of recurrent disease, the detection of limited metastatic spread does not necessarily provide information about tumor aggressiveness. Consequently, selecting an appropriate treatment strategy that balances potential clinical benefits against treatment-related risks remains challenging in patients with BCR⁶–⁸.

In a search for clues to better understand metastatic progression and to help tailor personalized care in PC we turned to the physical endurance of CTCs and to characteristics of their cancer-promoting cellular microenvironment. All live CTCs isolated from the peripheral blood already survived at least two journeys through the heart and were exposed to considerable fluid shear stress (FSS) harmful to non-adapted cells^1,2^. Therefore, it is expected that these CTCs present phenotypes supporting “mechanical fitness”. Cell’s adhesion, deformability (ability to change shape) and stiffness (resistance to change shape: an inherent ability of elastic deformation) constitute mechanical phenotype specific for each CTC. In our previous works we pointed at elevated adhesion as the major mechanical property distinguishing CTCs from tumor residing cells^3^. High adhesion supports CTCs clustering and clusters are acknowledged precursors of metastasis^4^. Clustering also protects CTCs from the direct attacks of macrophages. Moreover, we found that CTCs are pre-equipped with high adhesion by contacts with tumor associated macrophages (TAMs) that accompany them to the circulation as a part of protective CTC microenvironment^3^. In addition to high adhesion, we found before that elevated deformation, alongside of low stiffness, set apart castration resistant (disease progression despite castrate levels of testosterone) from castration sensitive patients and amount for “mechanical fitness” of CTCs^5^. CTCs are diverse, with a wide range of epithelial and mesenchymal characteristics, amounting to epithelial-mesenchymal plasticity (EMP)^3,6^. Moreover, a distinctive subset of CTCs: the tumor hybrid cells (THCs) that are products of incomplete phagocytosis of tumor cells by macrophages (MΦs), displays epithelial, stem cell and macrophage markers and their enumeration associates with aggressive disease^7,8^. The heterogeneity of CTCs was long considered a roadblock in studies on these cells and their use in biomarking. We aimed to determine whether the well-recognized heterogeneity of circulating tumor cells (CTCs), and more broadly of tumor-associated circulating cells (TACCs), reflects distinct strategies of mechanical adaptation to the hemodynamic and biomechanical stresses encountered during circulation, and whether these adaptive phenotypes have potential biomarker value. We found that distinct mechanical adaptation strategies may define functional subpopulations of CTCs and TACCs, potentially providing new biomarkers of tumor aggressiveness, metastatic potential, and disease progression.

## Materials and Methods

### Human subjects

Liquid biopsy blood samples were obtained by morning phlebotomy from male patients in the University of Texas Health Science Center at San Antonio/Mays Cancer Center and Audie L. Murphy Memorial VA Hospital in San Antonio, TX following the approved Institution Review Board protocols (IRB#HSC20130219H, CTRC#13–0001, Mays Cancer Center Biorepository Protocol). Written informed consent was obtained from all subjects in compliance with the Declaration of Helsinki, Belmont Report, US Common Rule following the US Department of Health and Human Services and the FDA regulations and Good Clinical Practice guidelines. Patients’ information is provided in **Tables 1-3**. Included in the analysis is data on blood samples collected for 23 patients that were subjects of our previously published work^3^. The clinical information such as time from liquid biopsy to progression was updated for these patients, and additional data not used before such as tumor hybrid cells enumeration, was included.

**Table 1.** Information on 77 cases of liquid biopsies from 62 prostate cancer patients. Listed times are in days.

| **Total number of cases: 77**<br>(62 patients; 15 patients with two liquid biopsies, 3-12 months apart) |
|---|
| Race:<br>White (non-Hispanic): 44<br>White (Hispanic): 18<br>Black: 15 |
| Time from diagnosis to liquid biopsy:<br>average: 1767<br>median: 1120 |
| Time from liquid biopsy to progression:<br>average: 821<br>median: 705 |
| Non-metastatic: 35<br>Metastatic: 42 (22 synchronous)<br>M1a: 13<br>M1b: 25<br>M1c: 4 |
| Castration sensitive: 37<br>non-metastatic: 24<br>metastatic: 13<br>Castration resistant: 40<br>non-metastatic: 15<br>metastatic: 25 |
| Patients in groups based on frequency of CTCs:<br>Average (A): 33<br>Deformable (B): 12<br>Highly adhesive (C): 17<br>Stiff (D): 15 |

### Prostate-Specific Membrane Antigen-Positron Emission Tomography/Computed Tomography (PSMA-PET/CT)

For selected patients (**Table 3**) whole body (skull base to mid-thigh) PSMA-PET/CT scans were collected with Siemens Biograph mCT20 PET CT close to the time of liquid biopsy (+/-one month; 14 cases), +/-2-4 months (8 cases) or no more than 9 months before the blood collection (2 cases). ^68^Ga-PSMA-11 was used in all but three cases (see **Table 3, patients #3, #11 and #17**). Quantitative analysis was performed with ePROMISE software to generate Maximum Standardized Uptake Value (SUVmax) of the most PSMA-avid metastatic lesion (SUVmax met) and SUVmax of prostatic bed (SUVmax local).

### Isolation and enumeration of cells from blood samples

Tumor associated circulating cells (TACCs) were isolated by microfiltration of blood using ScreenCell CC Filtration Kits (ScreenCell, Westford, MA)^9^. Cells collected on filters were subjected to nanomechanical phenotyping and stained with specific antibodies: anti-EpCAM-FITC or alternatively anti-vimentin-Alexa488, and with anti-CD163-PE and anti-CD80-Cy5. Brightfield and fluorescent images of cells on filters were used for enumeration of TACCs^9^. The following classes of TACCs were distinguished: epithelial-like canonical CTCs (EpCAM^+^ vimentin^+/-^CD80^−^CD163^−^), designated as “CTC-epi”, CTCs undergoing epithelial-to-mesenchymal transition (EMT; EpCAM^−^vimentin^+^CD80^−^CD163^−^; CTC-EMT). Macrophages (EpCAM^−^vimentin^−^; MΦs) were classified into M1-like (CD80^+^CD163^−^; M1), intermediate (CD80^+^CD163^+^; inter MΦ) and M2-like (CD80^−^CD163^+^; M2). Tumor hybrid cells (THCs) presented EpCAM^+^CD80^+/-^CD163^+^ phenotypes. All TACCs and their clusters were enumerated. CTC-epi, CTC-EMT and THCs were nanomechanically phenotyped as the potential seeds of metastasis (“CTCs”). Macrophages were occasionally phenotyped for comparison with CTCs.

### Nanomechanical phenotyping of cells

Cells on filters were scanned with a Catalyst atomic force microscope (AFM; Bruker; Billerica, MA) mounted on a Nikon Ti inverted epifluorescent microscope to obtain maps of mechanical properties derived from force curves recorded for thousands of probe-cell indentation events. The SCANASSIST-AIR probes (Bruker) with spring constant determined with thermal tuning moved vertically (cell’s indentation) and laterally (scanning) to collect maps of cell’s properties using Quantitative Nanomechanical Mapping (PF-QNM) AFM mode. Data in five separate channels: peak force error (PFE), height, adhesion, elasticity and deformation were collected simultaneously during a scan that lasted up to five minutes per cell. Data from the first two channels, alongside collected brightfield inverted microscope images, were useful to assess cells’ morphology and clustering status. Nanomechanical parameters of adhesion (units: picoNewtons, pN), stiffness (elasticity channel; kiloPascals, kPa) and deformation (nanometers, nm) were extracted from the scans with NanoScope Analysis software v.1.7 (Bruker), using mode values for individual cells. Adhesion is defined as the cell’s ability to adhere to uncoated, chemically inert silicon nitride probe. Stiffness is the ability to be fully reversibly deformed (elastic deformation). Deformation is the ability to be both reversibly and not immediately reversibly deformed upon preset non-damaging force (elastic and plastic deformations). Cells for interrogation were randomly selected on filters until 10-20 cells per sample were phenotyped. The procedures are described in detail in Osmulski et al.^9^.

### Statistical analysis

Unsupervised hierarchical cluster analysis using Ward method with corresponding heat maps and principle component analysis of the mechanical properties of the cells, and cell enumeration was performed using OriginPro 2025 with additional assistance from Statistica (TIBCO). Binary logistic regression was applied to predict odds of cases (OriginLab). Kolmogorov–Smirnov test was used to compare data distributions of two unmatched groups. Normal distribution was checked with Kolmogorov–Smirnov and Anderson–Darling tests. To test whether Pearson correlation coefficients were significantly different from 0, we applied the two-tailed t test. For comparison of groups with normally distributed data, the post hoc Tukey test was summoned. If the normality test failed, the nonparametric Kruskal–Wallis test was applied and the Dunn test was used for between groups comparison. When the assumption of equal SD was not met and group sizes were substantially different, the Brown–Forsythe test was called. Then, the Dunnett T3 test was used for post hoc analysis. Statistically significant differences between populations were assumed if P < 0.05. General descriptive statistics was completed with OriginPro 2025. Soft cluster membership analysis was based on Futschik et al.(2005) and Kumar et al. (2007)^10,11^.

## Results

### Classification based on mechanical phenotypes of CTCs reveals the presence of discrete categories of cells and distinct groups of patients discerned by abundance of these CTCs’ categories

In a search for mechanical signatures of CTCs and their putative links to the disease progression, we mechanically phenotyped 1277 cells isolated by microfiltration from 77 blood samples from 62 patients (**Table 1**). As demonstrated in **Fig. 1A, B**, the cells conform to four categories, three of them with a single mechanical property elevated representing the highly adhesive, deformable, or stiff cells. The fourth “average” category displayed no outstanding mechanical feature (**Fig. 1A, B**). We hypothesized that the identified categories represent specific strategies of mechanical adaptations to circulation conditions. We further hypothesized that exceptionally adhesive, deformable or stiff cells possess distinct potentials to start metastasis. Next, we set to test how abundant are the cells’ categories in patients (**Fig. 1C**). Unsupervised clustering of patients based on relative abundance of cells’ categories led to identification of four groups: with a very prominent contribution of “average” cells (from 15% to over 76%; group A), with up to 33% of highly deformable (group B) or highly adhesive (43%, group C) or exceptionally stiff cells (56% group D; **Fig. 1C**). Group membership did not seem to be associated with a stage of the disease at the time of liquid biopsy: in all the groups about a half of the patients have been already diagnosed with metastasis (48% to 59%) and close to a half (42% to 52%) had castration resistant cancer. However, trends emerged when the time to progression (TTP) post - liquid biopsy was analyzed. The differences were not statistically significant, nonetheless the average TTP in A-group was 994 days, in contrast with the average 622 days in C-adhesive group. The average TTPs post-biopsy in STI and DEF groups were reported as 735 and 678 days, respectively.

**Figure 1.**
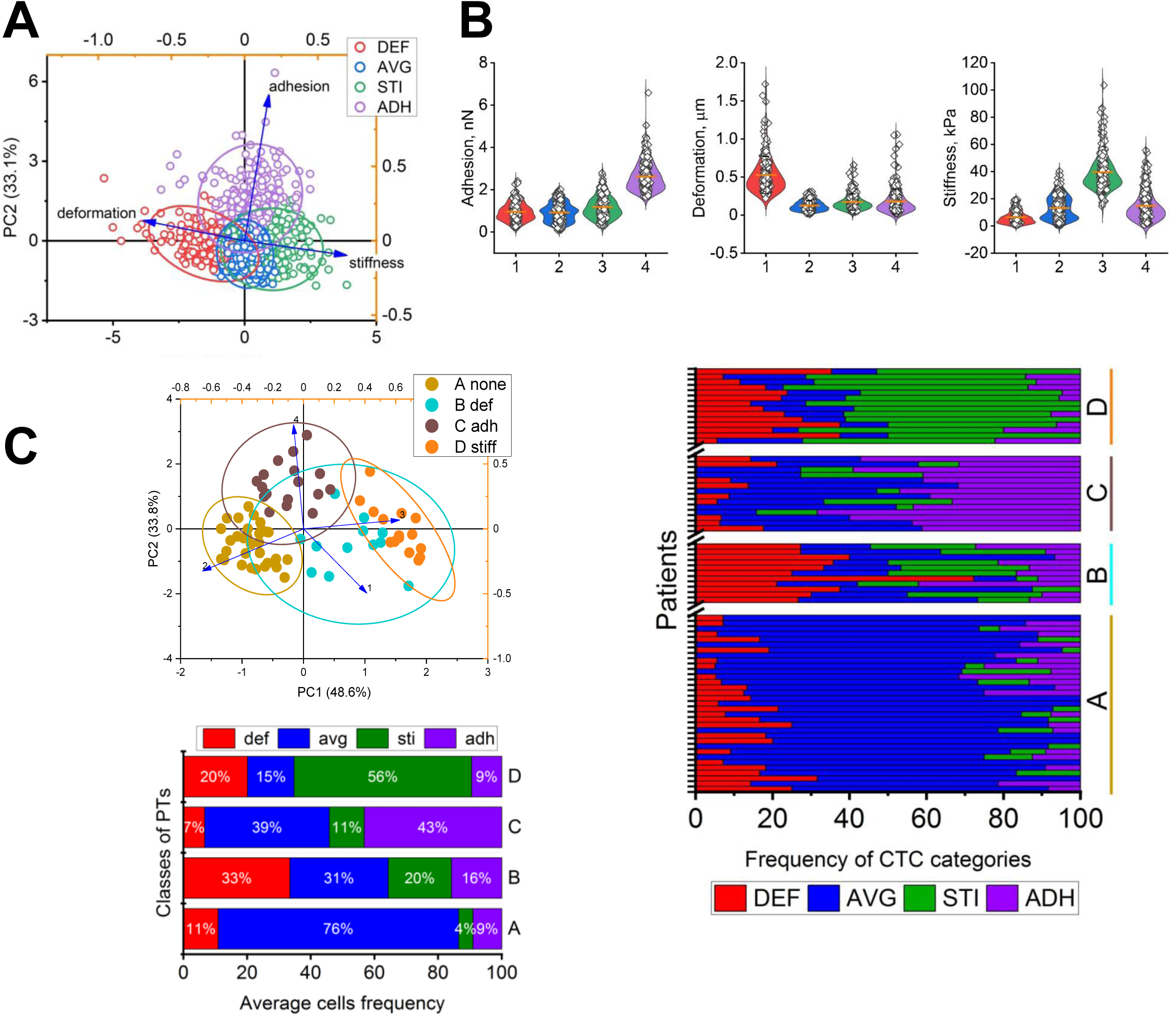
Mechanical phenotyping of CTCs isolated from 77 blood samples collected from 70 advanced (post-BCR) PC patients. **A.** Hierarchical cluster analysis of 1277 cells followed by principal component analysis (PCA) revealed the presence of four nanomechanically-defined categories: High-Adhesion (purple), High-Stiffness (green), High-Deformation (red), and Average (navy; no prominent mechanical parameters). **B.** Violin plots show mechanical parameters in each category. One dot = one CTC. **C.** Relative abundance of these CTC categories enables unsupervised clustering of patients into four distinct groups (A-D; *left*, one dot = one patient). The bar graph on the *right* presents the relative abundance of cells classified into the four categories: A, with most abundant cells from the “average” category, B-def, with prominent highly deformable cells, C-adh, with abundant highly adhesive cells and D-stiff, with abundant stiff cells. One bar = one patient. The bar graph in *lower left* presents average frequencies of cell categories in the A-C classes of patients.

### High adhesion of CTCs predicts early castration resistance

The large variations in post-biopsy TTP (**Table 1**) were not surprising taking into account the diversity of the patients’ cohort, with biochemical recurrence as the only selection measure, regardless the timing of BCR diagnosis. To limit the patients diversity, we focused on 30 patients starting first-round of androgen deprivation therapy (ADT) at the time of liquid biopsy, and followed clinically for at least a year to collect progression-free survival (PFS) time (**Table 2**). Cluster analysis of mechanical properties of CTCs from this sub-cohort resulted again in separation of cells into four categories, with 1-average, 2-stiff, 3-deformable or 4-highly adhesive cells (**Fig. 2A**), closely matching the categories defined for the large cohort (**Fig. 1A**). When the patients in sub-cohort were grouped by similarity of CTCs class frequency, the four groups of patients emerged, ascribing to cases with predominant deformable (A), average (B), adhesive (C) or stiff (D) cells (**Fig. 2B**). For majority of cases, the classification performed for the sub-cohort matched that performed for the full cohort. For example, all six cases from the group with prominent content of highly adhesive cells (**Fig. 2B**) were also classified in ADH group presented in **Fig. 1B**. Next, we used the four groups to perform the Kaplan Meier analysis with the Cox model estimator (**Fig. 2C, D**). Apparently, the survival function for patients from group C (highly adhesive CTCs) stood out with the shortest PFS (**Fig. 2C**) and the hazard ratio of 1.234 (**Fig. 2D**). We followed clinically these patients for up to 4 years; however 6 months is a clinical landmark time in ADT therapy, used for a short ADT course and for a check of PSA levels to assess the efficacy of therapy. ADT-undergoing patients with sharply lowered PSA level are considered castration sensitive. We estimated that if a patient starting first-time ADT has at least 6 CTCs (or more than 35% all probed CTCs) with the minimum adhesion of 1.66 nanoNewtons (nN), the probability of progressing into castration resistance within 6 months is 2.5 times higher than for other patients. Thus, the frequency of highly adhesive CTCs and the high-adhesion strategy emerged as the significant (p<0.004) predictive biomarker for early castration resistance.

**Figure 2.**
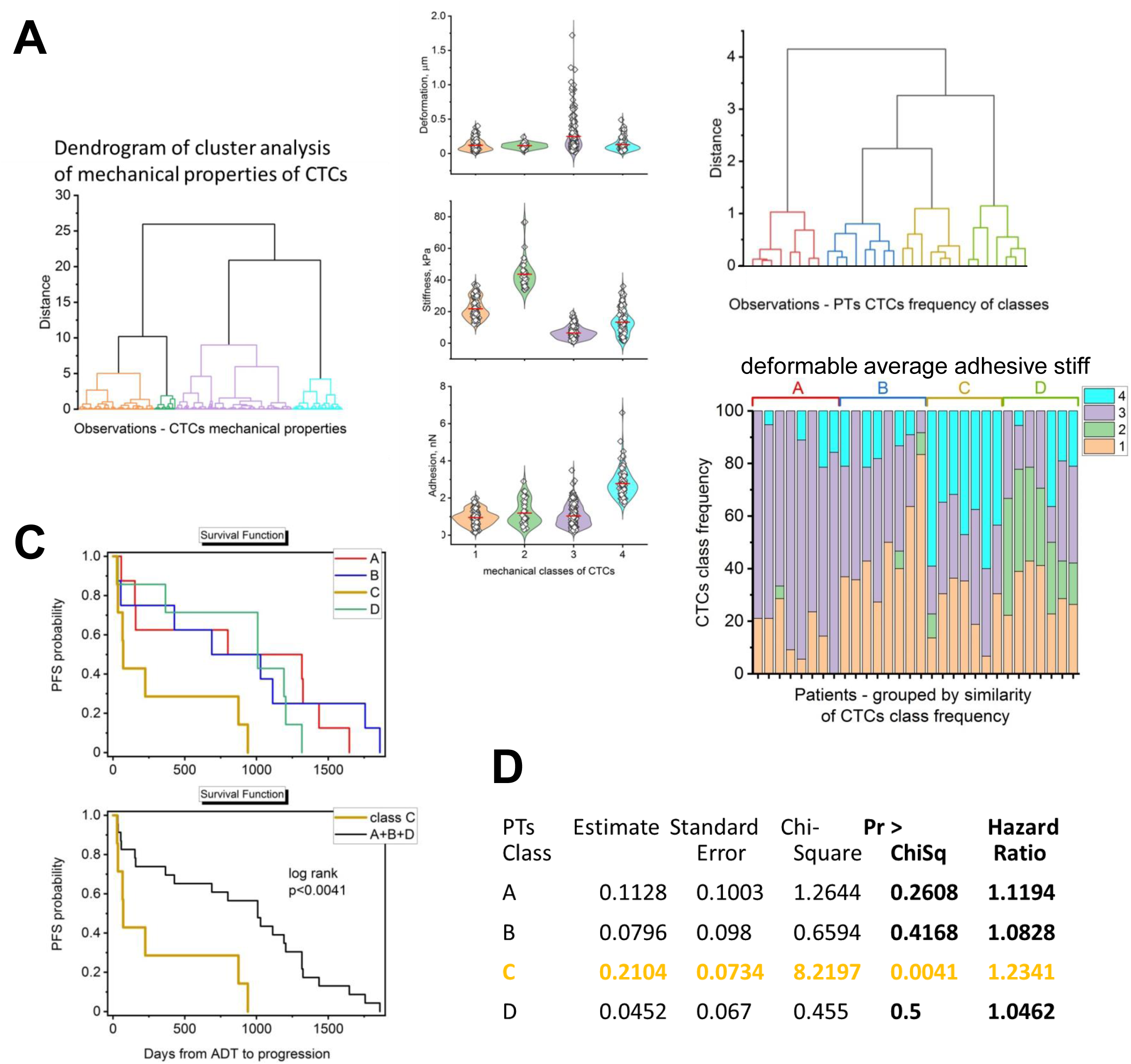
Nanomechanical phenotypes of CTCs isolated from 30 biochemically recurrent (BCR) patients starting first-round androgen deprivation therapy (ADT; see Table 2. **) A.** All CTCs from 30 PTs were grouped into 4 categories according to their mechanical parameters. Left: dendrogram, of cluster analysis, Right : violin plots demonstrating mechanical parameters of cells in each category. **B.** Relative abundance of these CTC categories enables unsupervised clustering of patients into four distinct groups (*top*). The column graph (*bottom*) demonstrates frequency of the CTC categories in individual patients, with A-D groups denoting similarity of the frequencies marked. One bar = one patient. **C.** Kaplan Meier estimator of progression free survival (PFS) probability based on enumeration of frequency of highly adhesive CTCs in BCR prostate cancer patients. *Left:* Data presented in B, with patients grouped by similarity of their CTCs nanomechanical phenotypes frequency into A-D classes, were used. *Right:* The A, B and D classes were pooled. The C class alone represented patients fast progressing toward hormone independence. **D.** Cox Model Estimator. Patients from class #C have a Hazard Ratio 1.234 (p<0.004) to fail ADT within the monitored period of over 5 years.

**Table 2.** Information on 30 BCR patients who started of 1^st^ round androgen deprivation therapy (ADT) close to the time of liquid biopsy, a subcohort of the 77 cases listed in Table 1. PFS - progression free survival from the time of ADT start or liquid biopsy. NA – not available.

| Sample Number | Disease spread at draw | PSA at draw (ng/ml) | Primary treatment | Stage at diagnosis, prostatectomy | Gleason: diagnosis, prosta-tectomy | Diagnosis to draw (days) | Draw to ADT (days) | PFS (days) | Group (Fig. 2C) |
|---|---|---|---|---|---|---|---|---|---|
| A1 | M1b | 125.98 | ADT | T2b, N0, M0, NA | 4+4, NA | 15 | 14 | 506 | A |
| A2 | none | 37.17 | radiation | cT3 | 4+3, NA | 540 | 0 | 152 | A |
| A3 | none | 1.36 | surgery | NA,<br>pT3a/b, R1, N0, Mx | 4+3, 4+3 | 596 | 35 | 122 | A |
| A4 | none | 7.1 | radiation | NA | NA | 2964 | -10 | 1647 | A |
| A5 | M1a | 0.64 | radiation | cT1c, N0, M0, NA | 3+4, NA | 1491 | 0 | 1436 | A |
| A6 | M1a | 0.05 | surgery | NA,<br>pT3b, N1, M0, R1 | 3+4, 4+4 | 126 | 0 | 1316 | A |
| A7 | M1b | 8.64 | surgery | cT2b,N0,M0<br>pT2c,N0,M0,R1 | 3+4, 4+3 | 1127 | 0 | 1324 | A |
| A8 | M1a | 43.8 | ADT | cT1c, N1, M0 | 4+5, NA | 23 | 28 | 85 | A |
| A9 | none | 14.99 | ADT | T2b, N0, M0, NA | 4+4, NA | 16 | 14 | 1874 | B |
| A10 | M1b | 704.3 | ADT | cT4, N1, M1<br>NA | 4+5, NA | 7 | 0 | 54 | B |
| A11 | regional | 2.94 | surgery | cT2b, N0, M0<br>pT3b, N1, MX | 4+5, 4+5 | 1148 | 0 | 1757 | B |
| A12 | M1a | 0.06 | surgery | cT1c, N0, M0<br>pT3b, N1, M0, R1 | 3+4, 4+4 | 287 | 0 | 1116 | B |
| A13 | M1a | 48.81 | ADT | cT3b, N1, M1a, NA | NA | 35 | 0 | 33 | B |
| A14 | none | 18.3 | radiation | T1c, Nx, Mx, NA | 3+3, NA | 3544 | 0 | 427 | B |
| A15 | regional | 0.94 | surgery | NA,<br>pT3b, N1 | 4+3, 4+4 | 443 | 0 | 1028 | B |
| A16 | M1b | 11.17 | radiation | cT1c, N0, M0, NA | 9, NA | 3325 | 15 | 702 | B |
| A17 | regional | 14.57 | surgery | cT2b, N1, M0<br>pT3b, N1, M0, R0 | 4+5, 4+5 | 218 | 0 | 1191 | D |
| A18 | none | 7.22 | radiation | NA | 3+4, NA | 835 | 0 | 1204 | D |
| A19 | none | 18.3 | radiation | T1c, Nx, Mx, NA | 3+3, NA | 2062 | 0 | 1008 | D |
| A20 | none | NA | radiation | NA | NA | 2153 | -90 | 917 | D |
| A21 | M1a | 0.05 | surgery | NA,<br>pT3b, N1, M0, R1 | 3+4, 4+4 | 126 | 0 | 1225 | D |
| A22 | regional | 102.74 | radiation | cT2b, N1, M0, NA | 4+3, NA | 35 | 42 | 77 | D |
| A23 | none | 31.62 | ADT | cT2b, N0, M0, NA | 3+4, NA | 14 | 0 | 365 | D |
| A24 | M1b | 21.22 | ADT | cT3b, N0, M1b, NA | NA | 31 | 0 | 66 | C |
| A25 | none | 0.78 | surgery | cT2b, N0, M0, pT3b, N0, M0, R1 | 4+4, 4+4 | 524 | 7 | 939 | C |
| A26 | regional | 0.17 | surgery | T1c, N0, M0, pT2, N1, M0, R1 | 4+3, 4+3 | 328 | 0 | 34 | C |
| A27 | none | 18.3 | radiation | cT1c, Nx, Mx, NA | 3+4, NA | 1086 | 0 | 874 | C |
| A28 | M1b | 44.27 | ADT | cT3a, NA | 4+5, NA | 51 | 0 | 28 | C |
| A29 | none | 10.04 | ADT | cT2, NA | NA | 15 | 0 | 224 | C |
| A30 | M1a | NA | ADT | cT3a, N1, M0 | 4+4, NA | 8 | 0 | 70 | C |

### Input of high-adhesion strategy may have a prognostic value in recurrent patients at the risk of metastatic spread

Encouraged by the predictive power of elevated CTC adhesion in patients starting ADT, we evaluated the significance of high-adhesion strategy in 25 BCR patients subjected to PSMA-PET/CT (Prostate-Specific Membrane Antigen-Positron Emission Tomography/Computed Tomography) imaging for assessment of cancer spread (**Fig. 3**). As indicated in **Table 3**, the spectrum of disease in these patients were classified into four categories based on the imaging results: 0 – no abnormal PSMA uptake detected (**Fig. 3A**); 1 – local recurrence in prostate; 2 – local recurrence beyond prostate in nearby lymph nodes and pelvic structures; 3 – oligometastasis with no more than five low-volume distant lesions^12^ (**Fig. 3B**); 4 – polymetastasis with advanced, high burden distant spread.

**Fig. 3.**
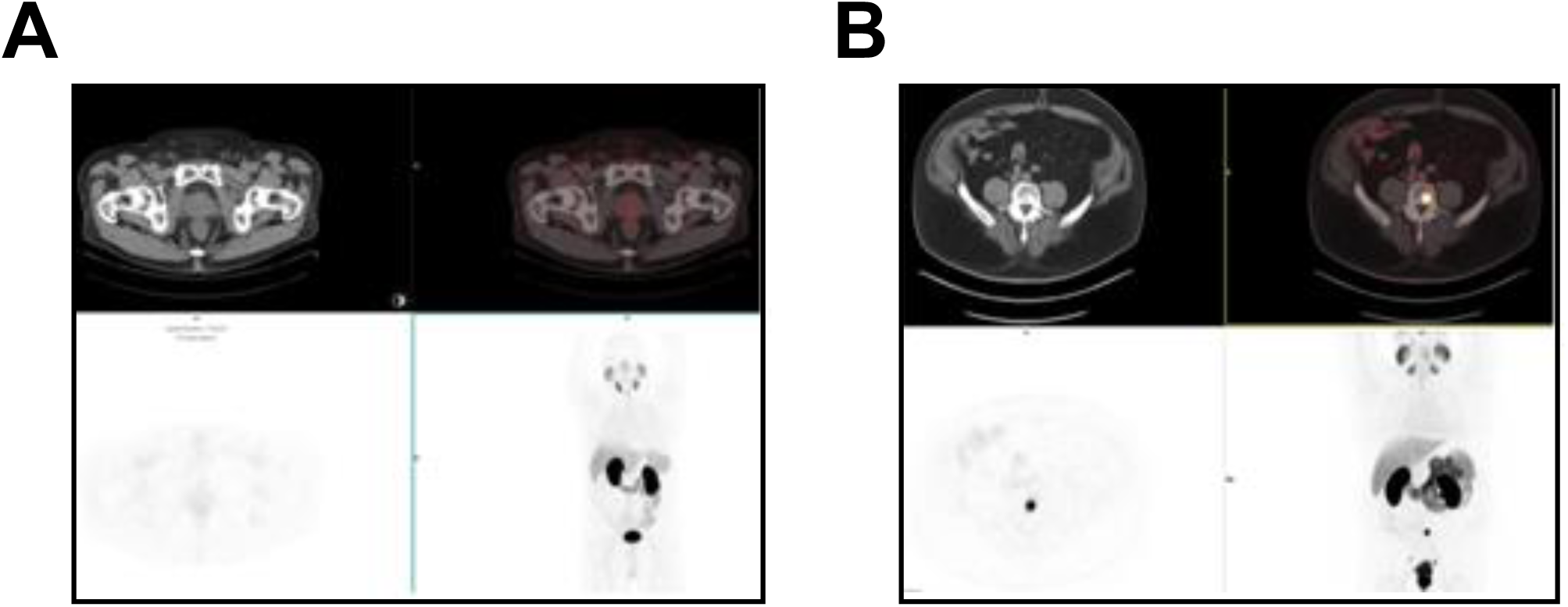
Representative PSMA-PET/CT images in ^68^Ga-PSMA-PET/CT scans of two recurrent patients. Axial CT and fused PET/CT (top images) and maximum-intensity-projection images (axial and whole-body) are shown for each category. **A:** Category 0, patient #3. No detectable PSMA uptake in prostatic bed, pelvic nodes or distant lesions. **B:** Category 3, patient #21. Oligometastasis with single bone lesion in L5 vertebral body. See **Table 3** for the patients’ information.

**Table 3.** Information on 24 patients who underwent liquid biopsy and PSMA-PET/CT scans for post-BCR prostate cancer. PSMA-PET/CT categories: 0 – no abnormal PSMA uptake detected; 1 – local recurrence in prostate; 2 – local recurrence beyond prostate in nearby lymph nodes and pelvic structures; 3 – oligometastasis with no more than five low-volume distant lesions; 4 – polymetastasis with high burden distant spread. SUV_max_: Maximum Standardized PSMA Uptake Value for local (prostatic bed) or distant (met) lesions. ⁶⁸Ga-PSMA-11 was used in all but three scans, with pifluostat-^18^F used for PTs #3, 11 and 17. L: Adhesion Score Low (<40% of mechanophenotyped cells with adhesion >1.66 nN), L: Adhesion Score High (≥40% of mechanophenotyped cells with adhesion ≥1.66 nN; HAS). LB - low burden, HD – high dilemma, HB – high burden. LN - lymph node. ND – no data. *Second visit for PT#1; a single scan was performed between the two visits.

| **Sample No.** | **PSMA scan category** | **SUV<sub>max</sub> local** | **SUV<sub>max</sub> met** | **PSA closest to draw (ng/ml)** | **Time from draw to scan (days)** | **Mean Adhesion [pN]** | **Group membership, comment** |
|---|---|---|---|---|---|---|---|
| 1 | 0 | 0.73 | 0.2 | 0.54 | 137 | 1287 | L, LB |
| 2* | 0 | 0.73* | 0.2* | ND | -97 | 1287 | L, LB |
| 3 | 0 | 2.03 | 2.03 | 6.21 | 6 | 1169 | L, LB |
| 4 | 0 | estimate | estimate | 0.2 | 59 | 1196 | H, LB |
| 5 | 0 | 0.17 | 0.51 | ND | -11 | 1570 | H, LB; 1Y earlier scan indeterminate for met |
| 6 | 0 | 1.37 | 0.78 | 12.2 | -99 | 963 | L, LB |
| 7 | 0 | 0.528 | 0.53 | 0.3 | -87 | 1767 | H, LB |
| 8 | 1 | 11.74 | 1.87 | 14.24 | 38 | 1247 | L, LB |
| 9 | 1 | 17.98 | 1.25 | 6.38 | 34 | 2879 | H, HD |
| 10 | 1 | 69.07 | 0.8 | 7.2 | -88 | 2208 | H, HD; 3<sup>rd</sup> BCR in 15 years |
| 11 | 1 | 17.51 | 1.43 | 18.45 | 6 | 928 | L, LB |
| 12 | 1 | 25.77 | 1.3 | 27.4 | -37 | 1650 | L, LB |
| 13 | 1 | 9.89 | 1.04 | ND | -148 | 1420 | L, LB |
| 14 | 2 | 12.85 | 17.49 | 1.53 | -56 | 1611 | H, HB; multiple local nodal; ADT in progress |
| 15 | 2 | 0.88 | 3.49 | 0.47 | 56 | 2528 | H, HD; indeterminate distant LN |
| 16 | 2 | 27.33 | 3.73 | 3.4 | -6 | 1280 | H, HB; ADT in progress |
| 17 | 2 | 12.19 | 15.37 | 82.8 | 3 | 1547 | H, HB; ADT in progress |
| 18 | 2 | 4.88 | 11.77 | 10 | -25 | 1378 | H, HB; multiple local nodal |
| 19 | 2 | 12.27 | 4.17 | 8.9 | 0 | 2067 | H, HD; suspicious external LN |
| 20 | 2 | 30.59 | 1.66 | 47.4 | 14 | 1855 | H, HD; suspicious external LN |
| 21 | 3 | 67.76 | 44.97 | 0.1 | -238 | 1908 | H, HB; single bone & LN; ADT in progress |
| 22 | 3 | 6.48 | 22.22 | 11.9 | -4 | 1489 | L, HB; post chemo for renal carcinoma |
| 23 | 3 | 2.93 | 1.9 | 7.2 | -31 | 4038 | H, HD; distant LN, bone in scan 3Y earlier |
| 24 | 4 | 1.27 | 115.29 | 11.91 | -281 | 2166 | H, HD; multiple bone |
| 25 | 4 | 5.97 | 59 | ND | 0 | 2849 | H, HD; multiple nodal, bone |

First, we tested if the border value of adhesion at about 1.7 nN computed from the Kaplan-Meier plot that signals a high risk of early castration resistance (**Fig. 2C**) is mounted in properties of CTCs beyond those from the ADT-starting patients’ subcohort. Peak deconvolution of the frequency distribution of adhesion values for 357 nanomechanically phenotyped single cells yielded four peaks: with maxima at 0.9 nN, 1.7 nN, 2.5 nN and the least pronounced fourth maximum at 4.7 nN (**Fig 4**). Apparently, the peak #2 at 1.7 nM matched closely to the proposed before border value of adhesion (1.66 nN, **Fig. 2**).

**Figure 4.**
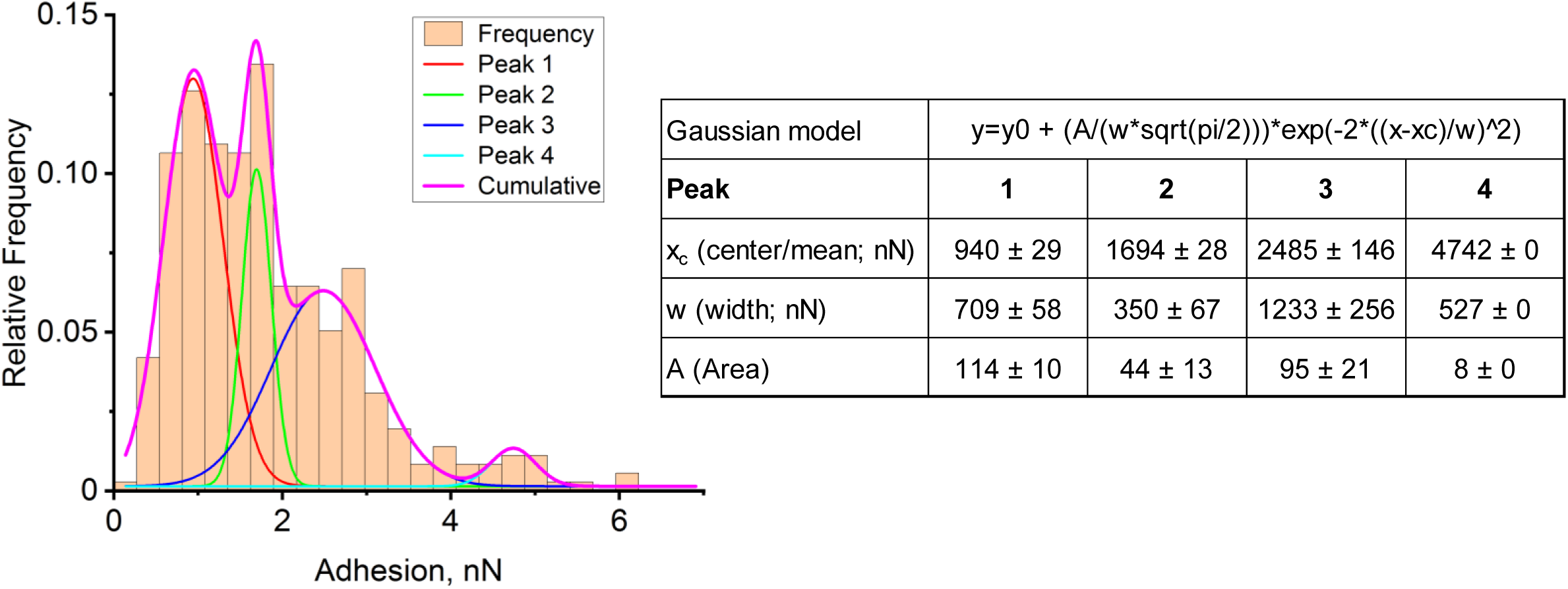
Frequency distribution of mechanical parameters of single CTCs isolated from the blood of prostate cancer patients. Histograms of relative frequencies for cell’s adhesion; Gaussian model peak parameters are listed on the right; reduced Chi-square of the model = 5.04E-05

We attempted to stratify patients based on frequency of their high-adhesion CTCs. Cells populating high-adhesion peaks #3 and #4 in the frequency histogram account for 40% of all cells (High Adhesion Score; HAS), as computed from the Area parameter of frequency histogram (**Fig. 4**). 9 patients belonged to low-adhesion group (Low Adhesion Score; LAS), consisted, with one exception, of categories 0 and 1, expected to pose a low concern for aggressive spread (**Table 3**). The sole exception was a category 3 patient with both renal and prostate cancers, post-chemotherapy, undergoing radiation and hormonal therapies at the time of liquid biopsy. The high-adhesion group consisted of 16 patients that had over 40% of their mechanophenotyped CTCs with adhesion higher than 1.66 nN. Not only metastatic patients (PSMA categories 3 and 4) were classified to this group, but also category 2 patients, who showed significant local spread and concerns for disease advancement. The most unexpected in H group, however, were three category 0 and two category 1 patients. We may suspect a false-negative scan result in patient #5 (category 0) since earlier scan of the same patient was concerning with indeterminate distant PSMA uptake. In turn, patient #10 (category 1) had a very long disease history (**Table 3**). The other cases may call for additional tests.

The significance of high adhesion was confirmed with a separate analysis that included SUV_max_ of metastatic lesions (SUV_max_ met) and average adhesion of all mechanophenotyped cells for each patient. Unsupervised Ward clustering followed by principal component analysis identified three groups: with low or high imaging burden and a high-dilemma phenotype with disproportionately elevated CTC adhesion relative to its lower imaging burden (**Fig. 5A, B**; **Table 3**). Soft-cluster membership analysis further supported the three phenotypes while revealing a continuum of patient-level membership. Low-burden (LB) patients (n=10) showed strong Cluster 1 membership, high-dilemma (HD) patients (n=6) predominantly showed Cluster 2 membership with greater overlap, and high-burden (HB) patients (n=8) predominantly showed Cluster 3 membership (**Fig. 5C**). Bootstrap consensus analysis (300 resamples) supported the overall cluster structure (**Fig. 5D**). All six HD patients were classified to the High Adhesion Score group, including two category 1 patients, #9 and #10 (**Table 3**). Interestingly, the unsupervised clustering analysis singled out three category 2 patients: #15, 19 and 20 as high dilemma. In all three cases scans were indeterminate or suspicious for distant lymph nodes spread. The category 3 patient #23 was also assigned an HD designation rather than more expected high burden; in this case there was a discrepancy in outcome of scans performed a year apart (**Table 3**). Apparently, unsupervised clustering confirmed the warning signs for two category 1 patients and helped to stratify category 2 and 3 patients into those with stable disease and those with high risk of progression.

**Figure 5.**
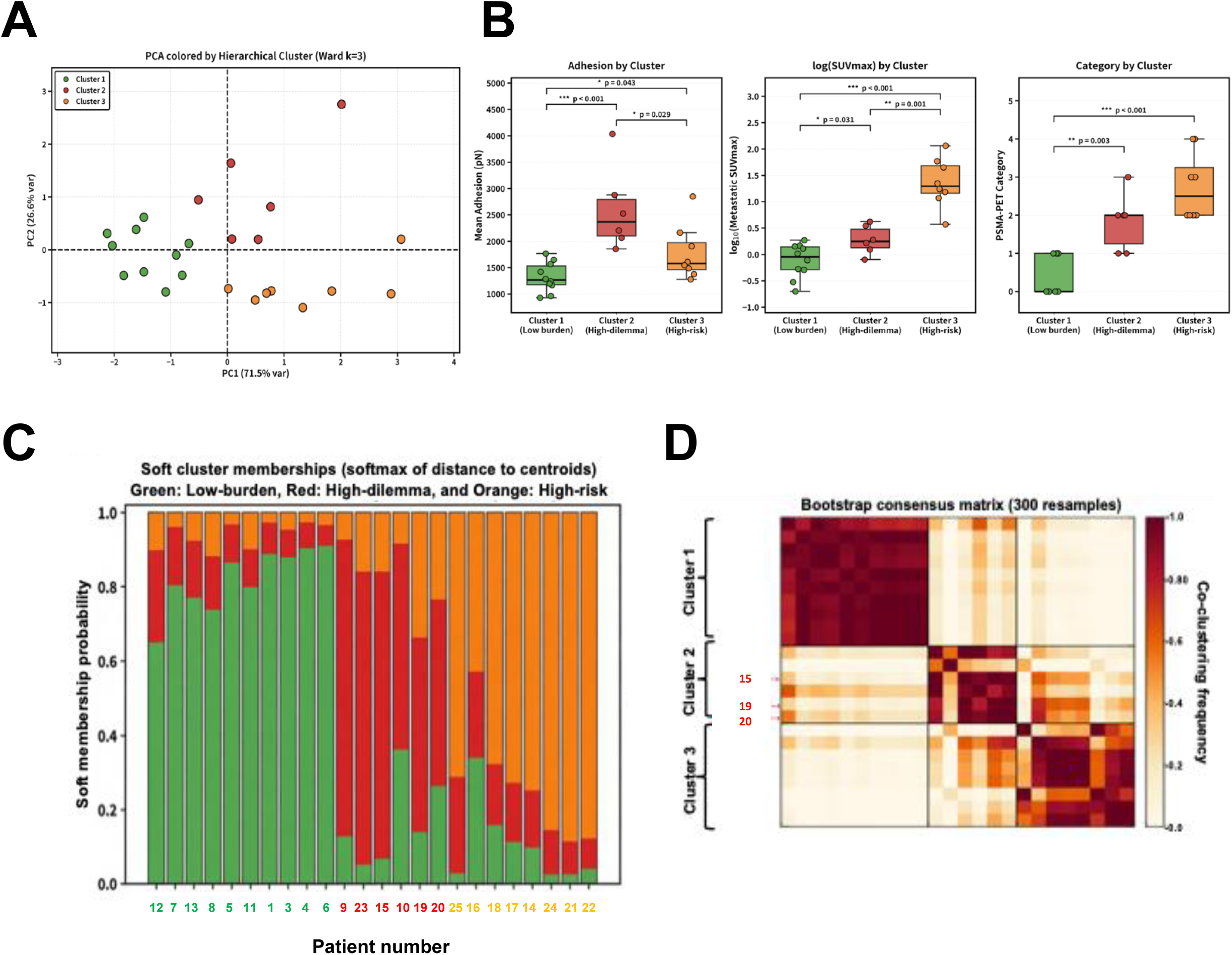
Classification of patients into “low burden”, high dilemma” and “high burden” groups based on average CTCs adhesion and outcome of PSMA-PET/CT scan. **A**, PCA of standardized mean adhesion force, log₁₀(metastatic SUVmax) and PSMA-PET category (n=24 patients) followed by unsupervised hierarchical clustering distinguished three clusters. Dots (patients) are colored in green (low burden, n=10) red (high dilemma, n=6) and orange (high burden, n-8). **B**, Box plots of the three input features stratified by cluster. Boxes show interquartile range, horizontal lines indicate medians, whiskers extend to 1.5 × IQR, with individual cases as points. Pairwise Mann–Whitney U tests were used for statistical comparison (*p < 0.05, **p < 0.01, ***p < 0.001). **C,** Soft membership probabilities were derived from distances to the three Ward cluster centroids. One column = one patient, with relative membership in the low-burden (green), high-dilemma (red), and high-burden (orange) phenotypes. **D,** Bootstrap stability of imaging – mechano CTC phenotypes. Bootstrap consensus matrix from 300 resamples shows the stability of the 3-cluster classification. Darker colors indicate more frequent co-clustering. PT#15, 19 and 20 (marked in red) show appreciable cross-cluster similarity between the high-dilemma (Cluster 2) and high-burden (Cluster 3) phenotypes.

### Stiffness and deformation related strategies provide additional tools for patients’ stratification

Next, we included additional parameters to the patient status analysis: SUV_max_ of prostatic bed (SUV_max_ local) and average stiffness of CTCs in addition to SUV_max_ met and average adhesion, while omitting the PSMA category to remove bias. PCA plot in **Fig. 6** demonstrates the unique positions of patients from PSMA category 0 (no detectable recurrence or spread) and categories 3 and 4 (distant spread): the former defined by stiff cells, the latter by highly adhesive cells and high SUVs. All but one category 1 cases were confined to the low-adhesion quadrant, with the exception of HAS and HD patient #9 (arrow in **Fig. 6**). Category 2 cases were diverse, with two of them worth to note. Patient #16 (HAS, HB) with ADT in progress, clustered together with category 1 cases in upper left quadrant. Another category 2 patient undergoing ADT, #17 (arrow in **Fig. 6**), clustered with other category 2 and 3 cases, from HAS and HB or HD groups. Both #16 and #17 were assigned HAS and HB membership. We may speculate that including stiffness as an additional mechanical parameter would help to further stratify BCR patients and to predict the response to therapy.

**Figure 6.**
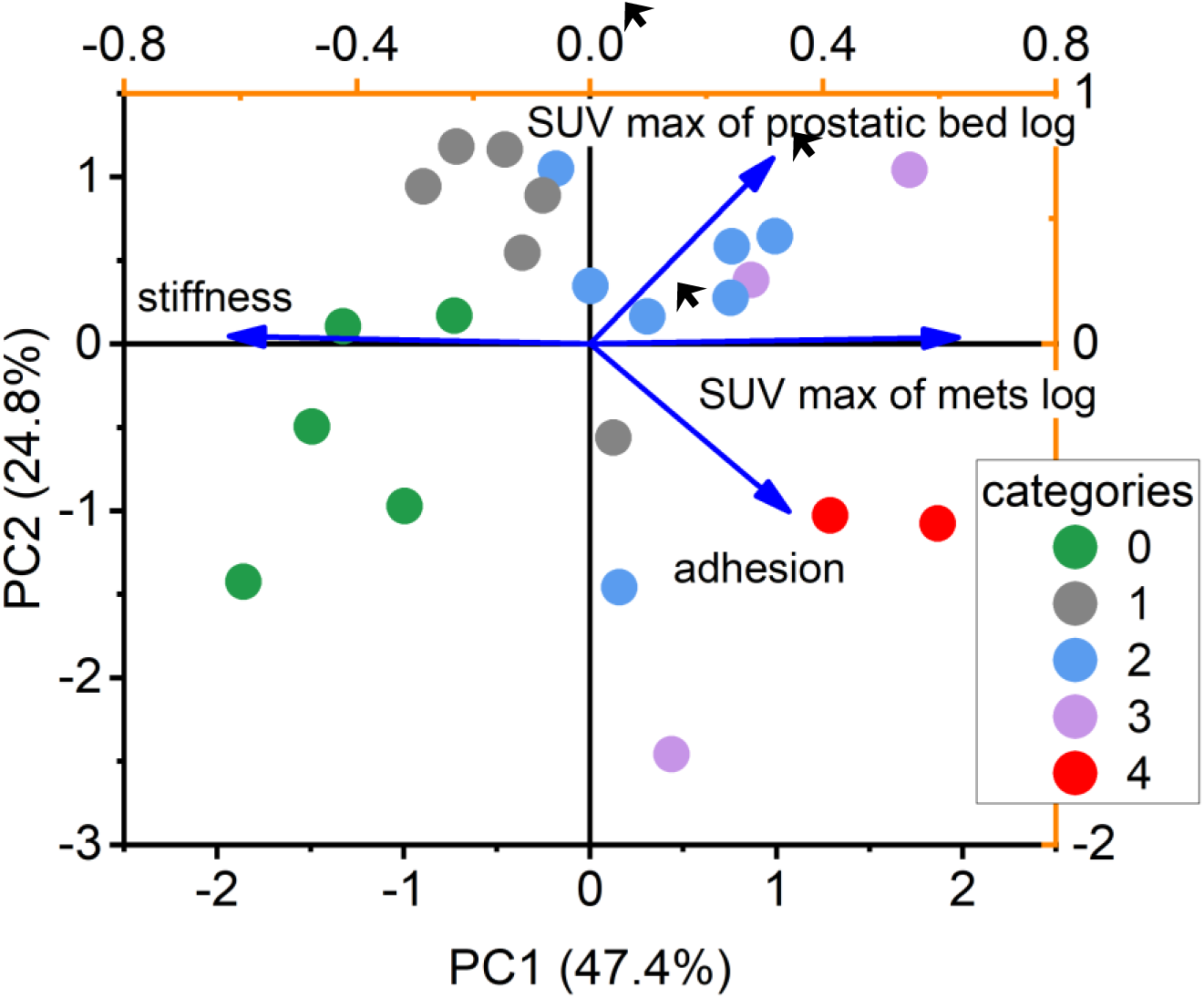
High adhesion of CTCs together with high uptake of PSMA defines metastatic patients. Principal component analysis of logs SUV_max_ of prostatic bed and of metastatic lesions, together with average stiffness and adhesion of mechanophenotyped CTCs. In a test analysis loading for average deformation was very similar to that of stiffness, therefore deformation was omitted from the demonstrated analysis. One dot = one patient, colored according to PSMA category. Arrows point at category 1 (grey dot) patient #9 and two category 2 (blue dots) patients #16 (left) and #17 (right); see text for descriptions.

In the above example higher average stiffness of mechanophenotyped CTCs’ was associated with low disease burden in categories 0 and 1, and with presumed better favorable response to therapy in category 2 patient #16; **Fig. 6**. The association of high stiffness of CTCs with low disease burden was also apparent in the patients starting 1^st^ round of ADT therapy listed in **Table 2**. In group D with abundant stiff cells (**Fig. 2B**) only two out of seven patients (29%) were metastatic at the time of ADT start. In all other groups at least half of the patients were metastatic (50% in AVG, 57% in ADH and 63% in DEF; **Table 2**).

### Properties of microenvironment of circulating tumor cells associate with mechanical adaptation strategies and add predictive value

Isolation of TACCs with the microfiltration method allows collecting TACCs defined as large cells including invasive CTCs and accompanying circulating tumor-associated macrophages. These cells often travel clustered and comprise the “CTC microenvironment”. In our previous work, we demonstrated associations of enumerations of distinct classes of TACCs with the prostate cancer burden^3^. For example, post-ADT patients had higher enumerations of macrophages with both M1-like and M2-like characteristics (inter MΦ) than pre-ADT patients with the less advanced disease^3^. Now, we found that patients with elevated frequency of cells belonging to high-adhesion strategy (group C, **Fig. 1C**) showed significantly higher enumeration of CTC-EMT than patients in other groups (**Fig. 7A**). In turn, the CTC microenvironment in patients with abundant deformable cells (group B; **Fig. 1C**) was characterized by high content of inter MΦ (**Fig. 7B**).

**Figure 7.**
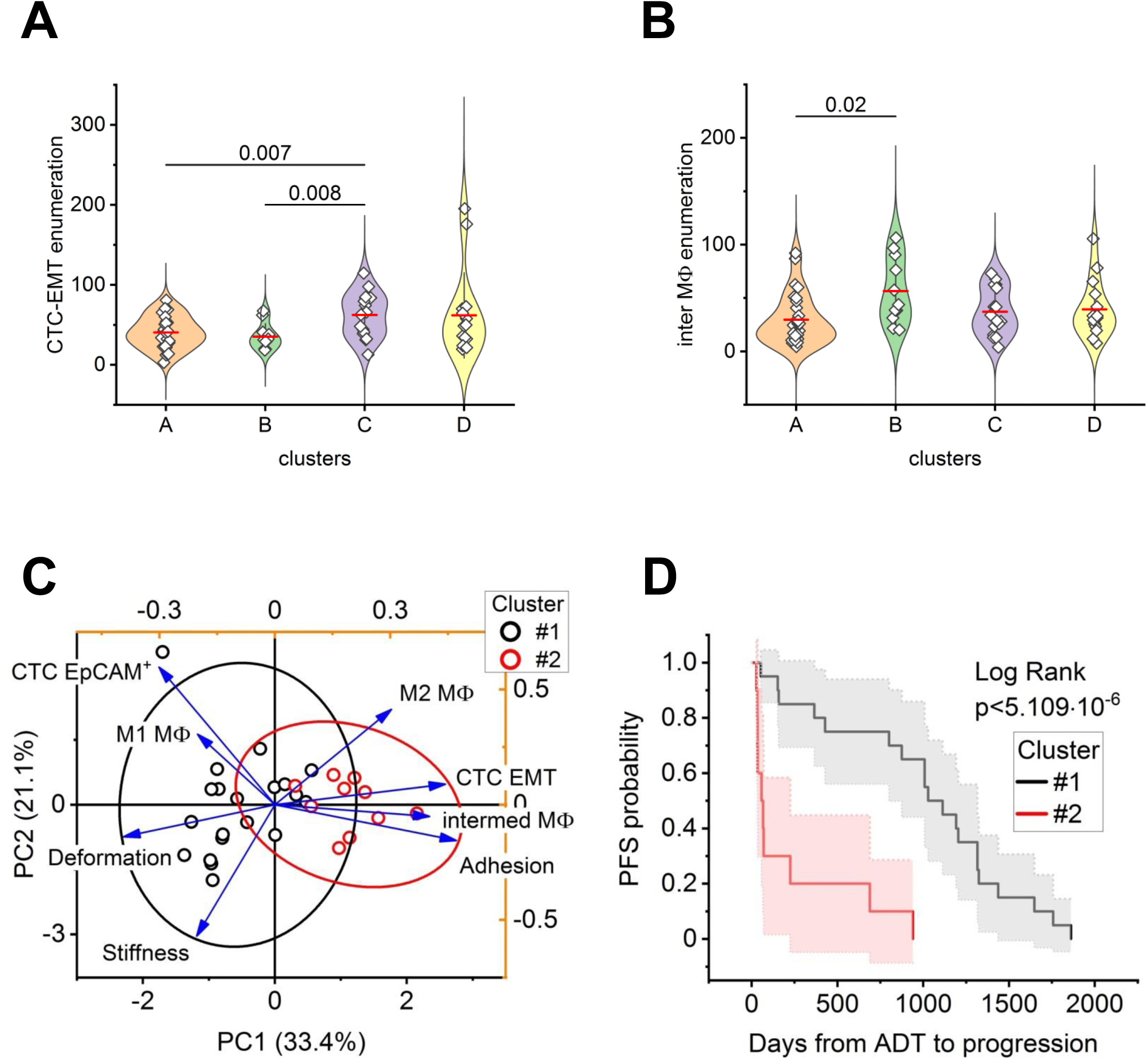
Enumerations of TACCs associate with mechanical adaptation strategies and help in prediction of early castration resistance. **A, B.** 77 phlebothomy samples were analyzed (**Table 1**), with patients separated into four groups based on frequency of CTCs belonging to distinct mechanical phenotype categories (**Fig. 1C**). High enumeration of EMT-undergoing CTCs (CTC-EMT) was a hallmark of patients from group/cluster C with abundant highly adhesive cells (**A**), whereas a high enumeration of inter MΦ distinguished group/cluster B from the “average” group A patients (**B**). **C,D.** Phlebothomy samples from 30 patients starting ADT were analyzed (**Table 2**). **C**. Cluster analysis of TACCs properties identified two groups of PTs. PCA (principal component analysis) groups PTs based on AFM derived mechanical parameters of CTCs and enumeration of TACCs. Patients with lower CTC deformation but higher adhesion and enumeration of EMT CTCs and intermediate MΦs showed significantly shorter time to progression (cluster #2) than PTs in cluster #1. **D.** Kaplan-Meier estimator shows highly statistically significant difference of PFS probability between 2 identified clusters of PTs. 80% of PTs from cluster #1 remained PF after 1 year vs. 19.6% from cluster #2. Per PT, ca. 20 CTCs were mechanically profiled with AFM and up to 300 TACCs were classified with immune stain.

To test the potential biomarking value of CTC microenvironment characteristics, we added enumeration of epithelial-like (EpCAM^+^; CTC-epi) and EMT-undergoing CTCs as well as distinct classes of MΦs to the nanomechanical phenotypes of the subcohort of 30 BCR patients starting their first round of ADT. Now, two clusters of patients were distinguished, with high adhesion in cluster #2 accompanied by high enumerations of CTC-EMT, inter MΦs and M2 macrophages (**Fig. 7C**). Kaplan-Meier estimator (**Fig. 7D**) confirmed the difference in PFS probability between the two clusters, with statistical significance at p< 5×10^−6^ stronger than that found for adhesion alone (p<0.004; **Fig. 2C**), with additional patients classified as the high-risk. Apparently, about 90% of patients classified in cluster #1 and less than 30% of patients from cluster #2 were expected to remain castration sensitive at the 6-months mark from the start of ADT.

### Tumor hybrid cells become exceptionally adhesive and abundant with disease progression

Next, we attempted to search if any specific kind of circulating invasive cells is responsible for the high adhesion phenotype. Three kinds of TACCs can potentially seed metastasis: canonical CTC-epi, CTC-EMT and tumor hybrid cells (THCs). CTC-EMT were an obvious choice due to their association with the ADH group of patients and with the risk of castration resistance (**Fig. 8A,C**). When enumeration of distinct TACCs was compared in patients with high (HAS) and low adhesion scores (LAS; **Table 3**), CTC-EMT were indeed significantly more abundant in HAS than in LAS (**Fig. 8A**). Interestingly, the fusions between immune cells, mostly macrophages, and tumor cells, THCs, were also significantly more abundant in HAS than in LAS (**Fig. 8B**). Enumeration of canonical epithelial-like CTC-epi did not differ between HAS and LAS (**Fig. 8C**). THCs constitute an important component of both the tumor and the CTCs microenvironment, and we found before that their content in blood increases with castration resistance^7^. In circulation, hybrids have the advantage of avoiding immune cells attacks by displaying “don’t-eat-me” signal and of metabolic plasticity with both cancer and immune cells features^7^. Moreover, THCs constituted about three-quarters of all enumerated invasive cells (76% in the whole cohort from **Table 1**), with non-significant differences between A-average (74%), B-deformable (83%), C-highly adhesive (75%) and D-stiff (76%) groups of patients distinguished in **Fig. 1C**. We set to explore the mechanical fitness of THCs isolated from the blood of patients listed in **Table 1**. As evident in **Fig. 8D**, in patients that were castration sensitive at the time of liquid biopsy THCs were less adhesive than a majority of CTC-epi and CTC-EMT. The pattern reversed in CR patients, with majority of THCs and only a few non-hybrid CTCs displaying exceptionally high adhesion (**Fig. 8E**). Such high adhesion was not observed in single cells, hybrids or non-hybrid CTCs, from CS cases (**Fig. 8F**). We concluded that the prominence of high-fitness, high-adhesion strategy of mechanical adaptation in advanced patients is supported mostly by THCs that change their mechanical phenotype toward higher adhesion with disease progression. Such change does occur in non-hybrid CTCs, however only to a small subset of cells (**Fig. 8E, F**). Since THCs comprise the majority of circulating invasive cells, we concluded that these cells are leaders in the high-adhesion strategy of mechanical adaptation.

**Figure 8.**
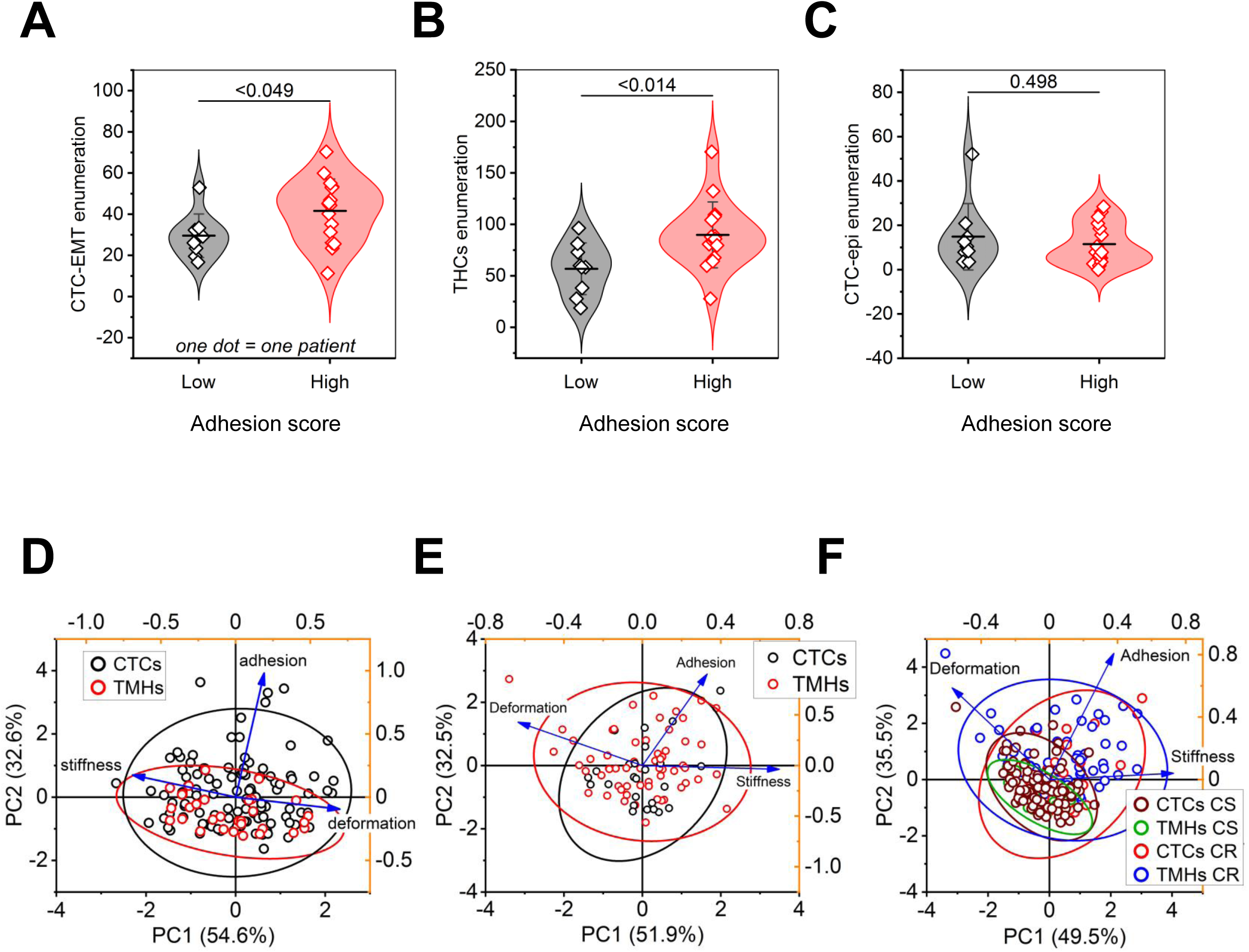
Tumor hybrid cells and CTC-EMT are abundant in patients belonging to high adhesion score group. These cells become more adhesive with the disease progression. **A-C.** Enumeration of TACCs was performed for the cohort of recurrent patients listed in **Table 3**. Among the invasive cells, CTC-EMT (**A**) and tumor hybrid cells (THCs; **B**) but not CTC-epi (**C**) were significantly more abundant in high adhesion score than in low adhesion score patients. One dot = one patient. **D-F**. Single cells capable of invasion (“CTCs”) from the cohort described in **Table 1** were mechanophenotyped and classified by surface markers. Among them, tumor hybrid cells (THCs, also known as tumor-macrophage hybrids; TMHs) were less adhesive than non-hybrid CTCs (labeled as CTCs) in castration sensitive patients (**D**), but more adhesive than CTC-epi and CTC-EMT in castration resistant patients (**E**). Generally, THCs in CR patients were the most adhesive among mechanophenotyped tumor associated circulating cells (**F**).

## Discussion

All living cells are adapted to their environment and the function that they carry out. The specificity of adaptations grows with the complexity of tissues and diversity of functions. Cells that need to rapidly change environments or are exposed to extreme conditions present especially intriguing examples of cellular plasticity. Circulating tumor cells (CTCs) shed from epithelial-origin tumors or metastatic lesions to the blood stand out among such cells. The contrast between the crowded solid tumor and often squeezed tumor cells, with supply of nutrients and abundant chaperone immune and fibroblast cells, and rapidly flowing fluid is drastic. Mechanical resilience of such cells holds the key to their survival and ultimate invasiveness^1^. Cancerous cells residing in tumors are known to be softer than their non-cancerous counterparts: a property useful in crowded environment with dense stroma^13,14^. If such cells are also poorly adherent upon gaining the mesenchymal-like phenotype, they may escape tumor via lymphatic vessels or invade extracellular matrix for the local cancer spread. CTCs with the very soft and non-adhesive phenotype, however, are vulnerable to damage by the fluid shear stress (FSS) and to cell death by anoikis^1^. This would be the most desirable fate for CTCs from a patient perspective. Unfortunately, live CTCs can be isolated from phlebotomy samples of advanced cancer patients even before the diagnosed distant spread, in numbers ranging from a few to several hundred per 7.5 ml of whole blood. All live CTCs pose the inherent danger of seeding metastatic lesions.

Here we focused on prostate cancer patients that experienced biochemical recurrence after a long period of dormancy, and/or were already diagnosed with local or distant spread. In our previous works we proposed that live CTCs isolated from peripheral blood are equipped with “mechanical fitness” phenotype to thrive in circulation^3,5^. Elevated adhesion bestowed by contacts with tumor-associated macrophages (TAMs) and supporting protective clustering was a key part of the mechanical fitness^3^. However, the widely acknowledged diversity of CTCs prompted us to explore mechanical parameters beyond adhesion. Indeed, the highly adhesive CTCs formed only a subset, about one-quarter, of over a thousand of mechanophenotyped CTCs, whereas highly deformable and very stiff cells fashioned other outstanding subsets. We propose that these cells represent diverse strategies of mechanical adaptations to circulation. Moreover, circulating tumor cell heterogeneity can be interpreted not as a predominantly molecular classification but as an integrated biological and biomechanical classification. To rationalize the identified strategies, we turned our attention to “professional” circulating cells such as erythrocytes and leukocytes. The reported stiffness of erythrocytes is remarkably similar to the “high stiffness” category of CTCs, ranging from several to over 40 kPa^15,16^. Therefore, we propose that the high-stiffness strategy represents erythrocyte-like adaptation to circulation. Mechanical phenotypes of leukocytes, in turn, indicate variable softness ranging from a fraction of kPa to less than 2 kPa^17^. Leukocytes are also deformable to enable phagocytosis^18^. In our set of CTCs the softest and most deformable are those in “high-deformation” category, thus designated as “leukocyte-like” adaptation. The most remarkable is, however, the “high-adhesion” category of CTCs. Adhesiveness of the professional blood cells is generally low, measured in tens of picoNewtons rather than in nanoNewtons, the latter being the case of CTCs^3,19,20^. Low adhesiveness helps blood cells to avoid being stacked in vessels. Invasive CTCs ultimately need to attach to the vessel walls for extravasation, however those isolated from liquid biopsies are in the circulation phase of their life-spans and elevated adhesion helps them to form abundant clusters that amount to protective cellular microenvironment. CTC clustering is an acknowledged phenomenon linked to cancer aggressiveness, with the problem of vessel blocking solved by a remarkable property of disassembly and re-assembly of clusters in capillaries^4,21^. Therefore, we consider the high-adhesion property of CTCs to bestow the “clustering” strategy of mechanical adaptation, ingenious for tumor-shed cells.

We found cells representing the strategies unevenly distributed in the patients. This enabled a search for associations between the mechanical properties of CTCs and the disease stage or progression. The prevalence of clustering strategy seemed to be especially important as a biomarker of CTCs’ aggressiveness. In a small cohort of patients starting 1^st^ round of ADT for recurrent or metastatic prostate cancer, assessing the abundance of highly adhesive CTCs enabled prediction of early castration resistance: a biomarking achievement not available by other measures. Moreover, we found that tracking highly adhesive cells may also add prognostic value to advanced diagnostic imaging. In recent years, PSMA-PET/CT scans entered the standard clinical practice for exceptionally sensitive detection of early spread in prostate cancer, albeit with limited prognostic capabilities^22–24^. However, analyzing the abundance of highly adhesive CTCs and establishing a “high adhesion score” for patients proved its value: it pointed at potential risk for patients with inconclusive scan data and identified “outlier” patients with low PSMA uptake (“clean” scans) but high content of potentially aggressive CTCs. Independently, unsupervised clustering of quantitative imaging data (SUV_max_) combined with average adhesion of patient’s mechanotyped CTCs allowed stratification of patients into three groups: with low disease burden, high-dilemma and high disease burden. The latter two groups presented high adhesion scores. High-dilemma patients may represent a transitional stage of active disease dissemination, in which highly adhesive CTCs remain abundant before substantial metastatic burden becomes radiographically apparent. Interestingly, adding average stiffness to analyzed adhesion and SUV_max_ data opened the possibility that admixture of cells representing erythrocyte-like strategy specifically indicated a positive outcome of therapy, an attractive notion requiring additional studies.

Since all mechanophenotyped CTCs, not only those representing the clustering strategy, have relatively high adhesion, it was not surprising that the microfiltration method of TACCs isolation always retrieved both clustered and single cells. We may suspect that some of the single cells were clustered in blood but become separated during TACCs isolation. Regardless the possibly transient clustering, enumeration of distinct species of immunostained TACCs revealed the value of CTC cellular microenvironment and potential biomarker significance of integrated analysis of mechanical phenotypes and microenvironment characteristics. The attempt to predict early castration resistance was much stronger when enumerations of circulating macrophages and individual enumerations of CTC-epi and CTC-EMT were added to mechanical phenotypes. It is ominous that the only clinically approved biomarking method involving CTCs relies exclusively on enumeration of EpCAM^+^ CTC-epi and neglects CTC-EMT and TACCs with immune cell markers^25^. In this and in our previous studies abundance of CTC-epi was associated with low-burden disease and with a positive outcome^3^. In contrast, the abundance of CTC-EMT associated with the clustering strategy. However, we found that tumor hybrid cells surpassed CTC-EMT both in their eminent adhesion and also in enumeration when following the disease progression. Apparently, mechanical phenotype was not constant for a particular cell type and was changing with disease progression; this was especially accurate for THCs but also for CTC-EMT. The changes parallel enrichment of CTC microenvironment with intermediate MΦs. It is tempting to speculate that contacts with MΦs in circulating clusters support high adhesion in THCs and CTC-EMT in a similar manner that TAMs preparation of the future CTCs in the tumor^3^ and formation of THCs from TAMs and CTCs in the tumor^7^. Most likely distinct species of CTCs do not automatically belong to a single strategy; they populate all of them. The concept of strategies of mechanical adaptations governing invasiveness of CTCs allows better understanding and pinpointing the dynamic nature of metastatic spread. The majority of patients possessed representations of all strategies. The partition of strategies abundance indicates the risk of progression. Even more important, the timing and extend of transitions between the strategies may be of a high and universal biomarker value.

## Acknowledgements

The work was supported by U01CA283749 (Huang), IMAT R61CA291213 (Osmulski & Gaczynska), S10OD034419-01 (Osmulski), DoD PC170821 (Huang), ThriveWell & Morrison Foundations (Gaczynska & Osmulski).

## Notes

### Competing Interest Statement

The authors have declared no competing interest.

